# Autonomously replicating linear plasmids facilitate the analysis of replication origin function in *Candida albicans*

**DOI:** 10.1101/551127

**Authors:** Swati Bijlani, Mathuravani A. Thevandavakkam, Hung-Ji Tsai, Judith Berman

## Abstract

The ability to generate autonomously replicating plasmids has been elusive in *Candida albicans*, a prevalent human fungal commensal and pathogen. Instead, plasmids generally integrate into the genome. Here, we assessed plasmid and transformant properties, including plasmid geometry, transformant colony size, four selectable markers, and potential origins of replication for their ability to drive autonomous plasmid maintenance. Importantly, linear plasmids with terminal telomere repeats yielded many more autonomous transformants than circular plasmids with the identical sequences.

Furthermore, we could distinguish by colony size, transient, autonomously replicating and chromosomally integrated transformants (tiny, medium and large, respectively). *Candida albicans URA3* and a heterologous marker, *ARG4,* yielded many transient transformants indicative of weak origin activity; replication of plasmid carrying heterologous *LEU2* marker was highly dependent upon the addition of a *bona fide* origin sequence. Several *bona fide* chromosomal origins, with an origin fragment of ~100 bp as well as a heterologous origin, *panARS*, from *Kluyveromyces lactis* drove autonomous replication, yielding moderate transformation efficiency and plasmid stability. Thus, *C. albicans* maintains linear plasmids that yield high transformation efficiency and are maintained autonomously in an origin-dependent manner.

**Importance:** Circular plasmids are important tools for molecular manipulation in model fungi such as baker’s yeast, yet, in *Candida albicans*, an important yeast pathogen of humans, prior studies were not able to generate circular plasmids that were autonomous (duplicated without inserting themselves into the chromosome). Here, we found that linearizing circular plasmids with sequences from telomeres, the chromosome ends, allows the plasmids to duplicate and segregate in *C. albicans.* We used this system to identify chromosomal sequences that facilitate the initiation of plasmid replication (origins) and to show that a ~100 bp fragment of a *C. albicans* origin, as well as an origin sequence from a distantly related yeast, can both function as origins in *C. albicans.* Thus, the requirements for plasmid geometry, but not necessarily for origin sequences, differ between *C. albicans* and baker’s yeast.

## Introduction

Plasmids are autonomously replicating extrachromosomal elements that facilitate molecular studies in bacteria as well as in yeasts and other fungi (1). Some yeast species carry natural plasmids, either as circles (e.g., 2µ in *Saccharomyces cerevisiae* (2)), or as linears (e.g., killer plasmids in *Kluveromyces lactis* (3), and mitochondrial plasmids in *Fusarium oxysporum* (4, 5)). Plasmid replication requires, among other components, an origin DNA sequence to which the origin recognition complex (ORC) binds. Origins of replication initiation (ORIs) on chromosomes and plasmids appear to have different sequence requirements in different yeast species (6). In *S. cerevisiae*, autonomously replicating sequences (ARSs: ORIs able to drive plasmid replication) are modular, requiring a minimum of 100 bp that includes an 11bp ARS consensus sequence (ACS) (7–9) and a T-rich ‘B element’ (10, 11). In most other organisms, the DNA requirements for centromere and ORI function are less well defined: *K. lactis* requires a 50 bp ACS that is necessary and sufficient for ARS activity (12); and *Schizosaccharomyces pombe* has no specific ARS consensus but requires a region of >500 bp with multiple A-T hook motifs that binds ORC (13–15).

In *Candida albicans,* a common human fungal commensal and an opportunistic pathogen, ORIs have been only partially characterized. *C. albicans* origins, like those of *S. pombe* and higher eukaryotes, have longer and less well-defined DNA motifs (16). Prior work with *S. cerevisiae* identified ARSs based on their high transformation efficiency (17, 18). Early studies found that *Sc*ARS plasmids with circular or linear geometry could be maintained autonomously for some time (19, 20). Work in *C. albicans* identified a few sequences that conferred high transformation efficiency on circular plasmids (21–27). However, the resulting transformants were either highly unstable (transient transformants) or the plasmid rapidly integrated into the genome (integrants). The *CaURA3* marker used in many of these studies, was later found to have an intrinsic weak ARS activity (28), and there was no direct evidence that replication initiated from the inserted sequences.

We previously used a machine learning approach to identify proposed-origins (pro-ORIs) based on ORC binding activity and nucleosome occupancy patterns (28). Four pro-ORIs were shown to be *bona fide* origins that produced replication bubble structures on non-denaturing 2-dimensional (2-D) DNA gels, thereby providing direct evidence of ORI function (28). Importantly, all four *bona fide* ORIs also drove plasmid replication on linear (but not circular) plasmids derived from circles carrying long inverted telomere repeats separated by a spacer sequence that is cleaved to linearize the plasmid (29). These large plasmids with inverted telomere sequences could work well, but were prone to rearrangement during propagation of the circular precursor plasmid in *E. coli*.

Here, we compared circular and linear plasmids in *C. albicans* that rely on *bona fide* ORIs for their maintenance. Linear plasmids were constructed from circles *de novo* by PCR with primers bearing telomeric repeats prior to transformation. Linear plasmids consistently had higher transformation efficiency, larger numbers of autonomous transformants and higher mitotic stability than analogous circular plasmids. Transformant colony size was a clear reflection of plasmid stability, with tiny colonies indicative of unstable, transient transformants, medium colonies indicative of autonomous transformants with moderate stability levels, and large, smooth colonies were indicative of integrants, in which plasmid was inserted at chromosomal positions. We also tested four markers, including *CaURA3* and *CaHIS1,* as well as heterologous markers, *CdARG4* and *CmLEU2* (30), which all had different levels of origin-dependent transformation efficiency and maintenance. Finally, we tested *bona fide* ORIs (28) as well as origin fragments and heterologous origin sequences, and found that a ~100 bp ORI fragment, and a *K. lactis panARS* (31) have moderate origin activity in *C. albicans*.

## Results

### Circular *CaURA3* plasmids with and without ORIs

Overall, across the markers and plasmids tested, three types of transformant colonies were evident. Tiny colonies that could not be maintained on selection (Fig. S1A and B) with undetectable plasmid retention (MS ~0) indicative of rapid plasmid loss were defined as **transient transformants** (referred to as **transients** hereafter) (Table 1). Large, round colonies, with short lag time and doubling time (Table 1, Fig. S1C) were defined as **integrants** based on their highly stability under selection (MS ~80-100%). Medium colonies (MS 1-80%) that grow, albeit less well than integrants under selection, with comparatively longer lag and doubling times (Table 1, Fig. S1D), were defined as **autonomously replicating transformants (ARS-transformants**) assuming that replicating plasmids can be maintained under selection, and lost in the absence of selection. Accordingly, colony size can reliably predict the MS of plasmids and used as a proxy for the number of different transformant types.

**Table 1.** Properties of different types of transformants obtained with circular and linear plasmids.

| Type of colony |  | Size (mm in diameter) | Lag time (min) | Doubling time (min) | Obtained with plasmids | Mitotic stability |
| --- | --- | --- | --- | --- | --- | --- |
| Transient | Tiny | $\leq 0.4$ | $174 \pm 5$ | $856 \pm 3$ | pCir/pLin-CaURA3 (±ORI410),<br>pCir/pLin-CaHIS1 (±ORI410),<br>pCir/pLin-CmLEU2 (±ORI410),<br>pCir/pLin-CdARG4 (±ORI410) | 0% |
| Autonomous | Medium | 0.4 – 1.6 | $30 \pm 10$ | $140 \pm 50$ | pCir-CaHIS1 (±ORI410),<br>pCir-CdARG4 (±ORI410),<br>pCir-CmLEU2 (+ORI410) | $\leq 10\%$ |
| | Medium | 0.4 – 1.6 | $28 \pm 6$ | $42 \pm 27$ | pLin-CaURA3 (±ORI410),<br>pLin-CaHIS1 (±ORI410),<br>**pLin-CdARG4 (±ORI410),**<br>**pLin-CmLEU2 (±ORI410)** | 10-40% |
| Integrant | Large | 1.6 – 2.2 | $19 \pm 2$ | $25 \pm 3$ | pCir/pLin-CaHIS1 (±ORI410) | 80-100% |

To test the hypothesis that *bona fide* ORIs drive the autonomous replication of plasmids in *C. albicans*, we first constructed circular plasmids with the *CaURA3* marker similar to those from prior studies (22, 23, 25–27) with and without *bona fide ORI410* (28) (Fig. 1A). We compared transformation parameters including transformation efficiency (TE, number of transformants/µg of DNA), size of the transformant colonies (tiny (<0.4 mm); medium (0.4-1.6mm) and large (>1.6mm), Table 1), mitotic stability (MS, proportion of cells that retain the plasmid under selection) and plasmid loss rate (LR, rate of plasmid loss per generation in the absence of selection). TE with and without *ORI410* was relatively modest (17 and 9 transformants/µg DNA, respectively (Fig. 1B)). Importantly, all selected colonies were tiny (<0.4 mm), with and without inclusion of *ORI410* (Fig. 1B). The tiny colonies did not grow upon re-streaking, or when seeded into liquid cultures (Fig. S1A), that is a characteristic of transients. Thus, as in several prior studies (22, 23, 25, 27), circular *CaURA3* plasmids were not maintained autonomously.

**Figure 1.**
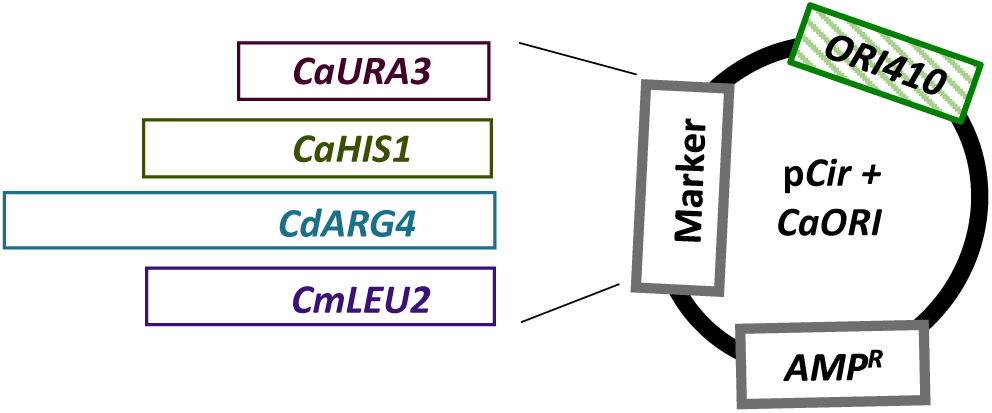
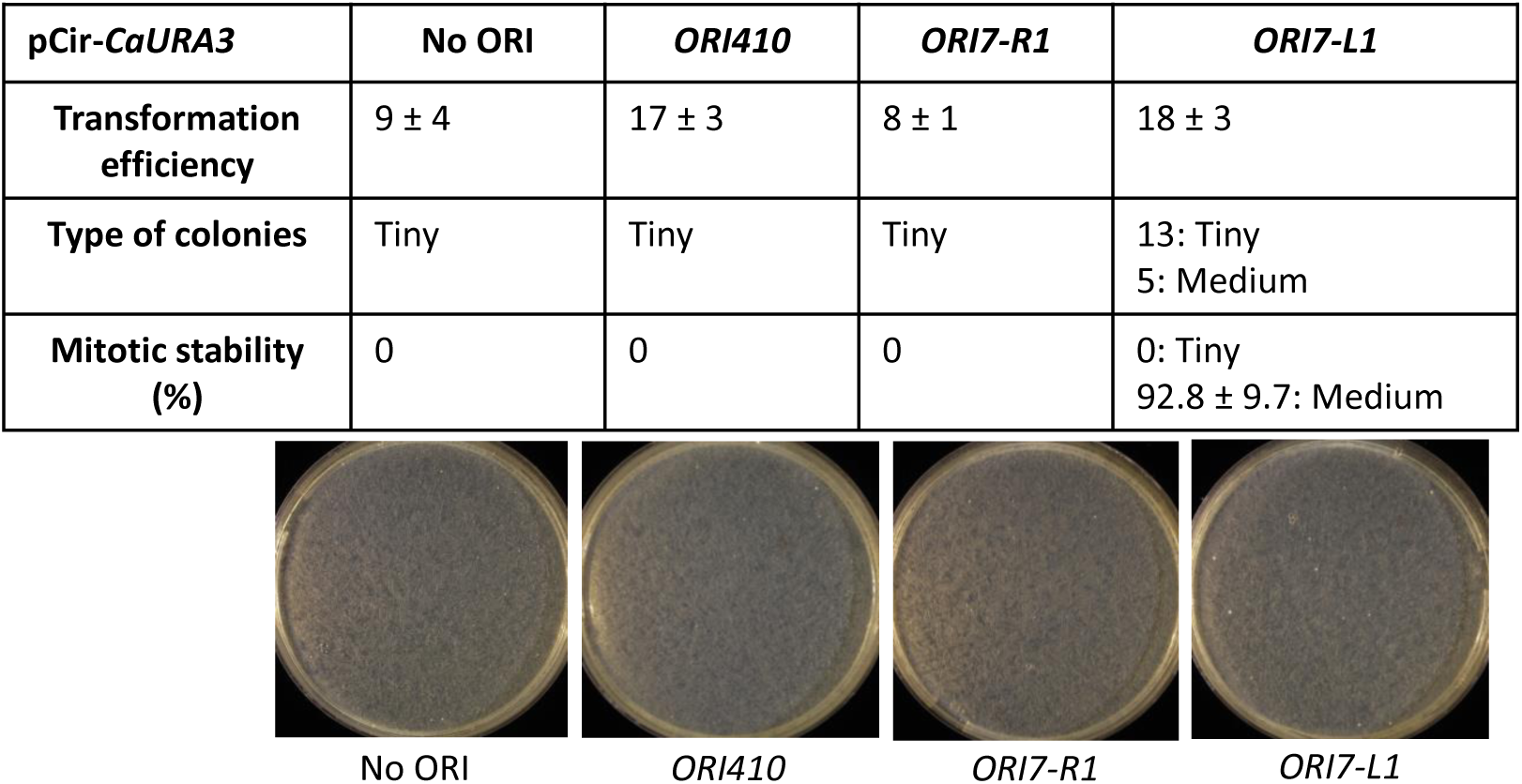
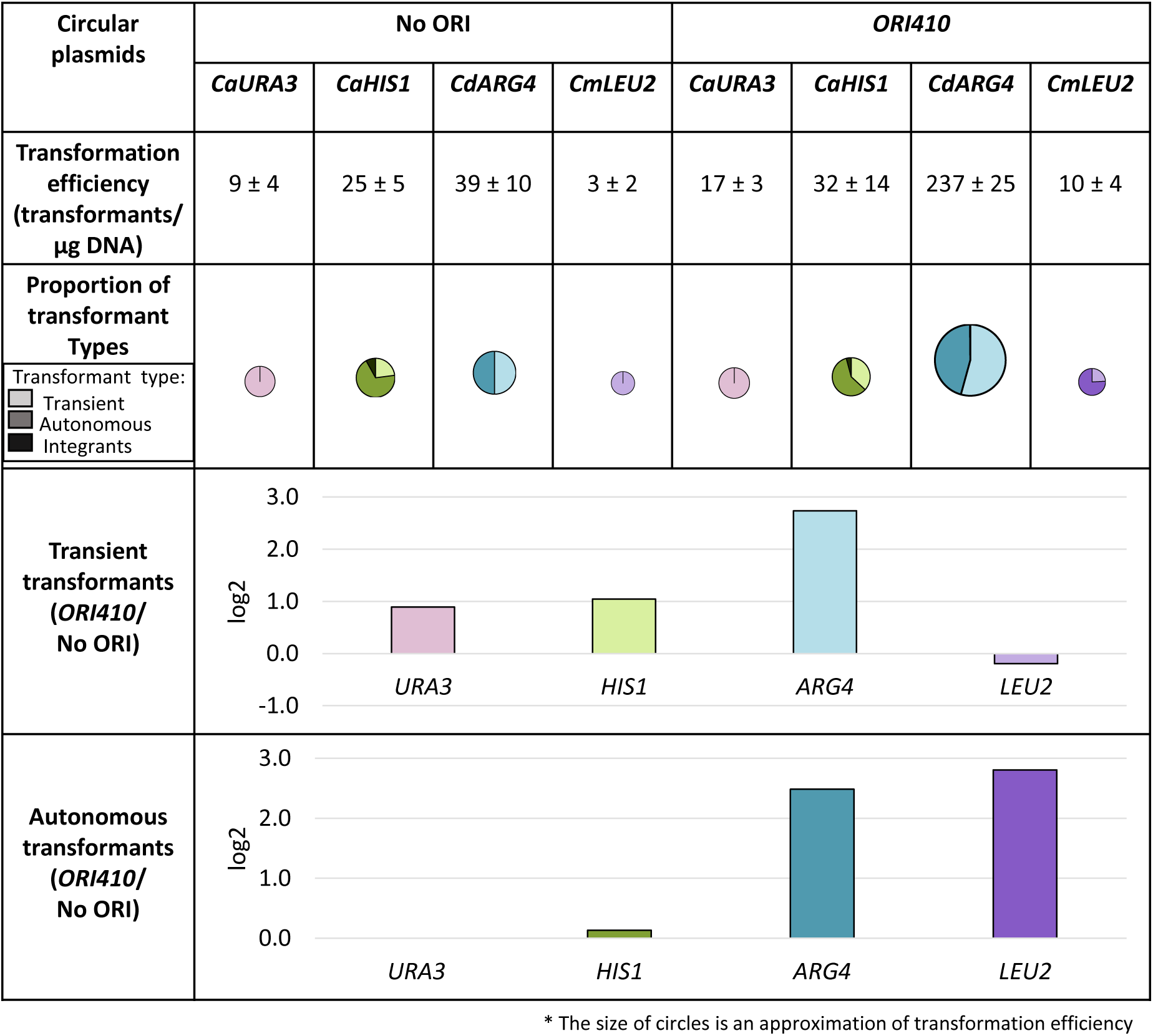
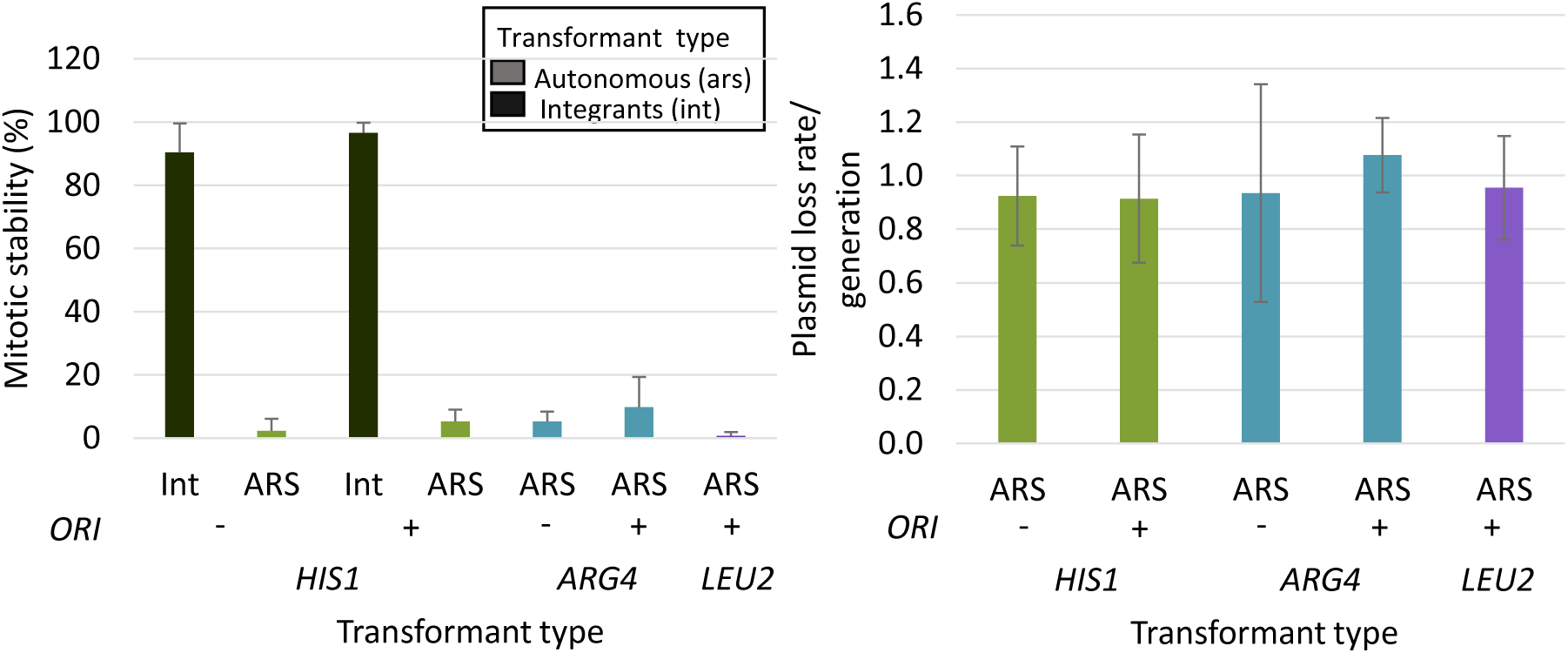
**(A)** Map of a circular plasmid showing relative position of selection markers (*CaURA3*, *CaHIS1*, *CdARG4*, *CmLEU2*) and ORI sequence. **(B)** pCir-*CaURA3* plasmid with different origin sequences: transformation efficiency, types of colonies and their mitotic stability. The transformation efficiency is an average of three independent experiments. **(C)** Comparison of circular plasmids carrying different selection markers with and without *ORI410*: transformation efficiency, proportion of different types of transformants and log2 value of the ratio of average number of transient or autonomous transformants with ORI to without ORI (*ORI410*/ORI^-^). Different markers are represented by different colors and different types of transformants are represented by varying shades of a color (lightest shade representing transients, intermediate shade representing ARS-transformants and darkest shade representing integrants). The transformation efficiency is an average of three independent experiments. **(D)** Mitotic stability (%) of integrants and ARS-transformants and plasmid loss rate/ generation for ARS-transformants obtained with different circular plasmids with and without *ORI410*. The data represents the average of three independent colonies of each type. Int: integrants; ARS; ARS-transformants.

Because this result conflicts with the claim that two sequences, *ORI7-R1* and *ORI7-L1*, drive the replication of a circular *CaURA3* plasmid (26), we constructed plasmids with these sequences in pCir-*CaURA3.* Both of them had modest TE (8 and 18 transformants, respectively, Fig. 1B). We obtained only transients for *ORI7-R1* with TE similar to the no-ORI plasmid; *ORI7-L1* gave twice as many transients compared to no-ORI plasmid, and produced a small number of stable transformants (Fig. 1B), indicating that they integrated into the genome. Thus, neither of the two *CEN7* flanking sequences yielded autonomous transformants in the context of a circular *CaURA3* plasmid (Fig. 1B), consistent with the poor performance of pCir-*CaURA3-ORI410*. Similar results for pCir-*CaURA3* with *ORI410*, *ORI7-L1* and *ORI7-R1* were also evident in a second strain background (Table S1).

### Comparison of different selectable markers on circular plasmid

We next asked if the *C. albicans HIS1* (*CaHIS1)* marker would show better TE and MS than *CaURA3*, with the goal of obtaining ARS-transformants. However, *CaHIS1* plasmid yielded small numbers of transformants (32 and 25, with and without *ORI410*, respectively), with a modest increase (~25%) in TE attributable to *ORI410* (Fig. 1C). pCir-*CaHIS1* ARS-transformants had MS <5% and plasmid loss rates of ~0.9 (Fig. 1D), indicating that they were autonomous but highly unstable. Thus, in addition to transients and integrants (analyzed in more detail below), pCir-*CaHIS1* produced a small number of ARS-transformants-a group not detected with pCir-*CaURA3* (Fig. 1C).

Because heterologous markers are less likely to integrate into the genome, we tested *CmLEU2* marker from *Candida maltosa* and *CdARG4* marker from *Candida dubliniensis* (30). With the addition of *ORI410*, TE of *CmLEU2* was increased by ~3-times (Fig. 1C), and most of them were ARS-transformants with MS <5% (Fig. 1D) compared to only transients without *ORI410*; no large colonies were detected. Thus, *CmLEU2* produced a small number of ARS-transformants with low MS upon addition of *ORI410*.

By contrast, *CdARG4* had a 5-fold higher TE with *ORI410* on the plasmid relative to without the ORI; ~50% being ARS-transformants (Fig. 1C) that had MS ~10% for those with *ORI410* and MS ~5% for those without the ORI (Fig. 1D). Thus, *CdARG4* with *ORI410* yielded more than 100 ARS-transformants/µg of DNA, with an improved MS (but with LR remaining quite high (Fig. 1D)). However, while *ORI410* was required for high TE, and improved MS, it was not required for some autonomous plasmid replication. We suggest that *CdARG4* sequence might enable a weak ORI to form on the plasmid (discussed later). Thus, for circular plasmids with all four selectable markers tested, the inclusion of an origin was not sufficient to produce relatively stable autonomously replicating plasmids (low MS and high LR). This indicates that a heterologous marker can drive autonomous replication of a circular plasmid with rare integration events, but they are lost at high frequency.

We also asked if autonomous plasmids were detectable in DNA extracts from the medium colonies (low MS and high LR). Indeed, Southern blot of DNA from medium colonies (Fig. S2A) detected bands with similar electrophoretic mobility to that of naked circular plasmids.

By contrast, in the DNA from a pCir-*CdARG4*-*ORI410* large colony with high MS (presumed integrant), a larger band was detected along with autonomously replicating plasmid (Fig. S2A), indicating integration in some cells in a population. This is consistent with the idea that large colonies contain integrated plasmid and medium colonies contain autonomously replicating plasmids. Moreover, analysis of the *CaHIS1* integrants found gene replacement at the native locus by single cross-over (Fig. S3A).

### Linear plasmids with telomere ends are maintained autonomously

Since circular plasmids did not yield high TE and high MS for ARS-transformants, we constructed and transformed linear plasmids, which are known to replicate autonomously in some fungal model organisms (4, 32–35). Since classical methods of producing linear plasmids (29) used for monitoring origin function in *C. albicans* (28) proved challenging, we designed a new approach in which linear plasmids were constructed from circular plasmids by PCR (details in methods, Fig. 2A) and transformed directly into *C. albicans.* Because telomere sequences are not necessary to be added to linear DNA during transformation in some fungal models (4, 32–34), we asked if the presence and the length of the telomere repeats (34 nt vs 57 nt TEL, i.e. 1.5X vs 2.5X of a single 23 nt *C. albicans* TEL repeat (36)) affects transformation parameters. Linear plasmids without TEL repeats had a TE of ~300/µg for all three markers tested (*CaHIS1*, *CdARG4* and *CmLEU2*) with the majority being transients (Fig. 2B). *CaHIS1* linear plasmid without telomere repeats, resulted in higher TE, increased number of ARS-transformants but also an increase in integration events compared to the corresponding circular plasmid (Fig. 2B vs Fig. 1C). Notably, *CmLEU2* linear plasmid without telomere repeats, resulted in a much higher TE and ARS-transformants (~50/µg DNA) than the corresponding circular plasmid (<10/µg DNA, (Fig. 2B vs Fig. 1C)). The *CdARG4* plasmid, was an exception yielding similar TE in circular and linear plasmid without telomeres (Fig. 2B vs Fig. 1C). However, all of the ARS-transformants obtained had low MS (<5%) (Fig. 2C) with an irregular colony shape, indicating that they were not readily maintained in the autonomous state and higher proportions of cells failed to divide in the colony under selection conditions.

**Figure 2.**
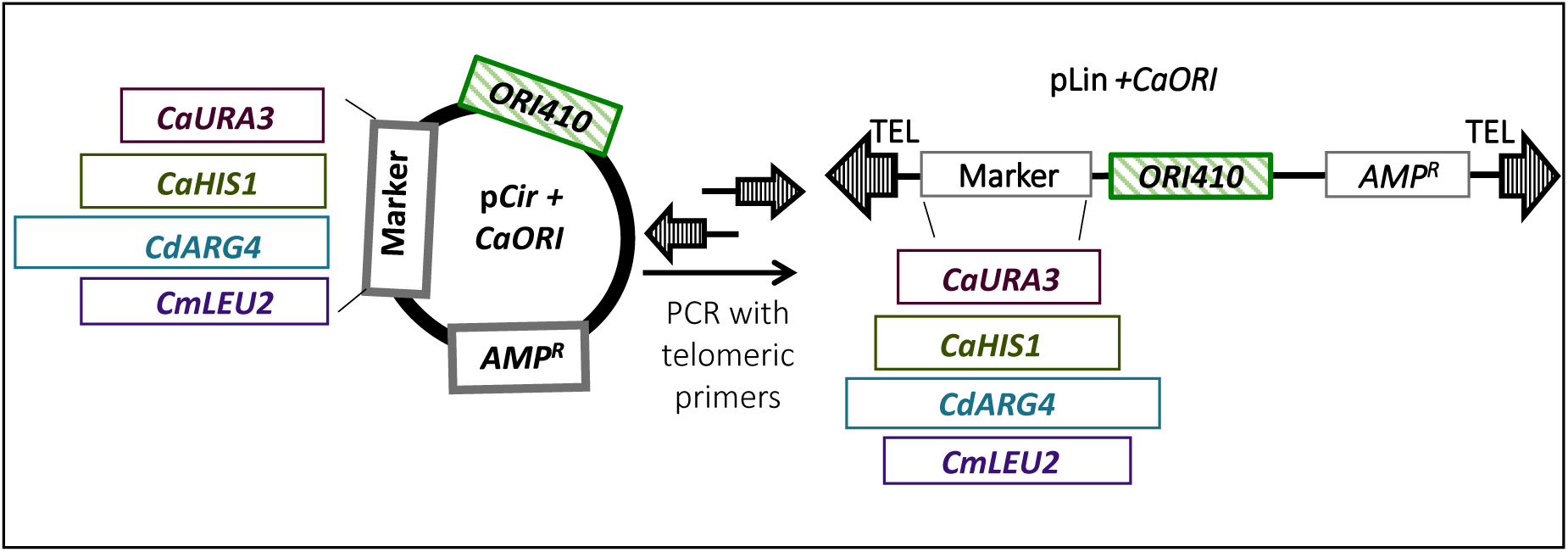
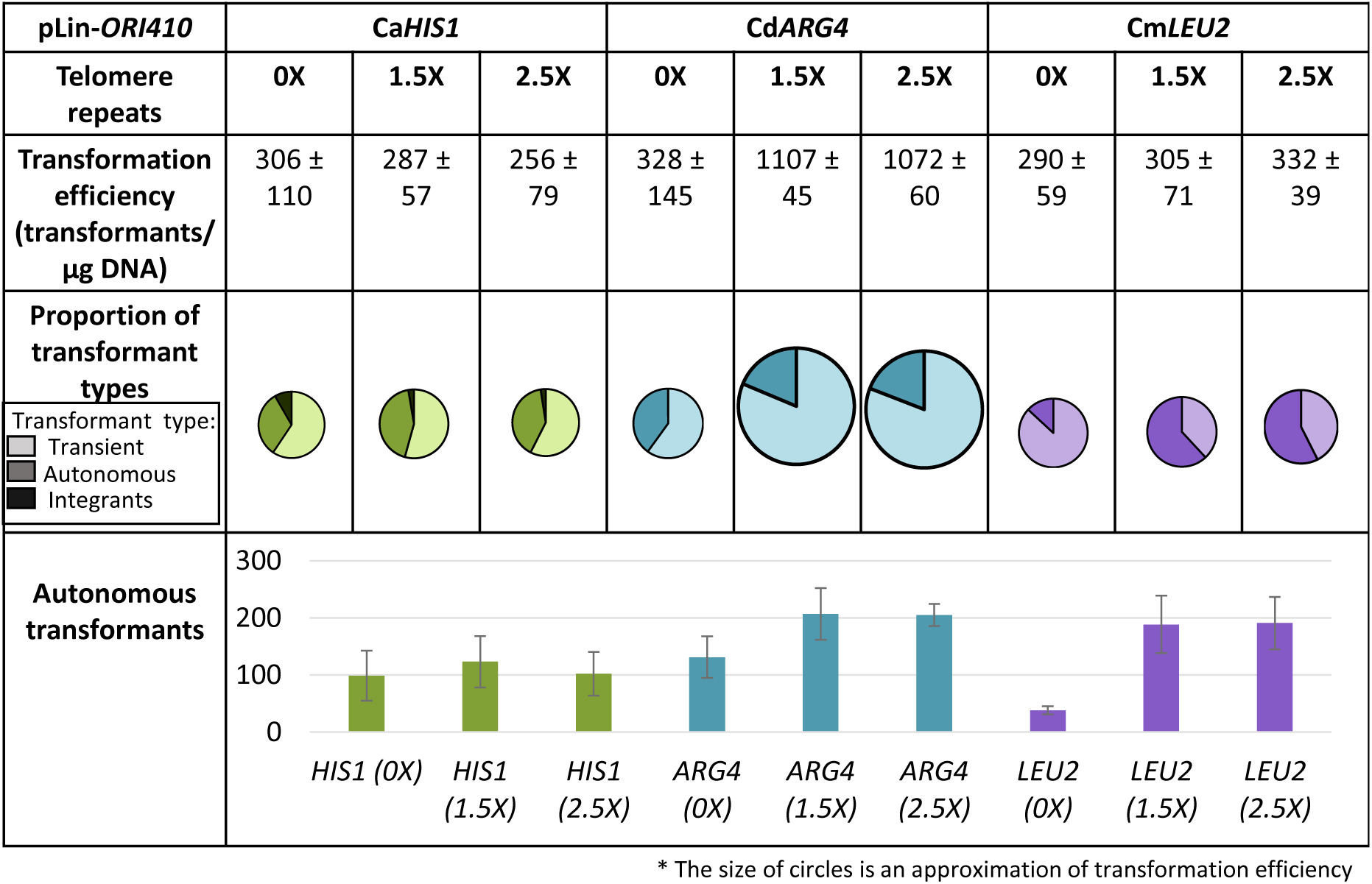
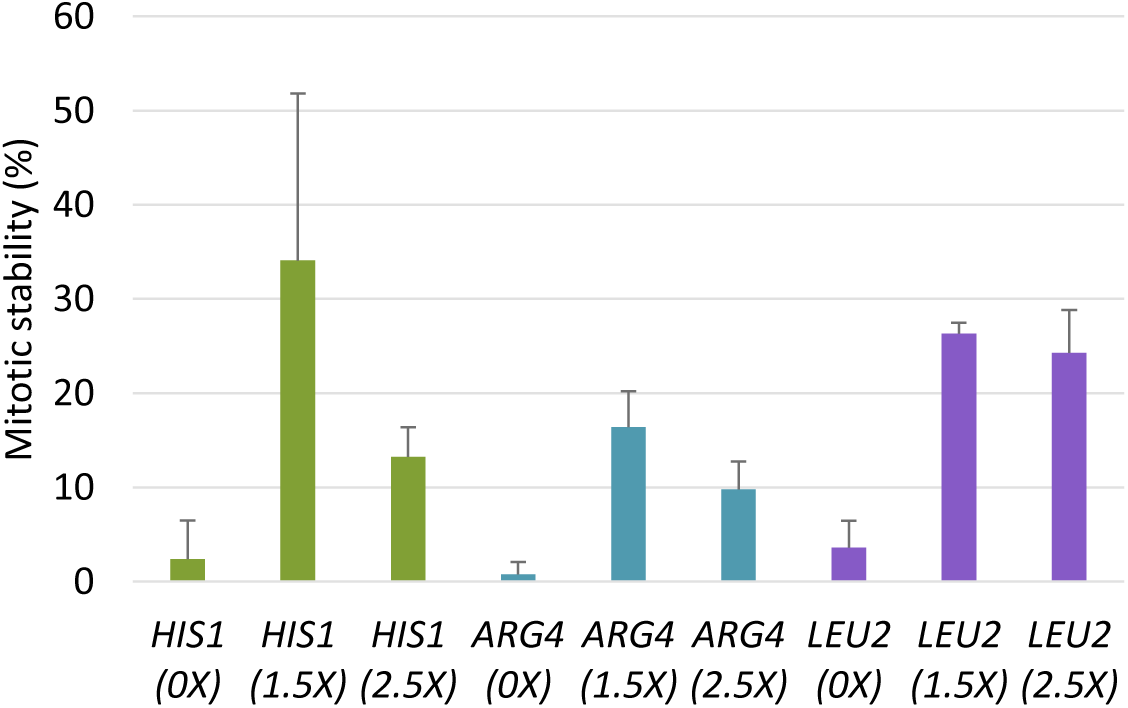
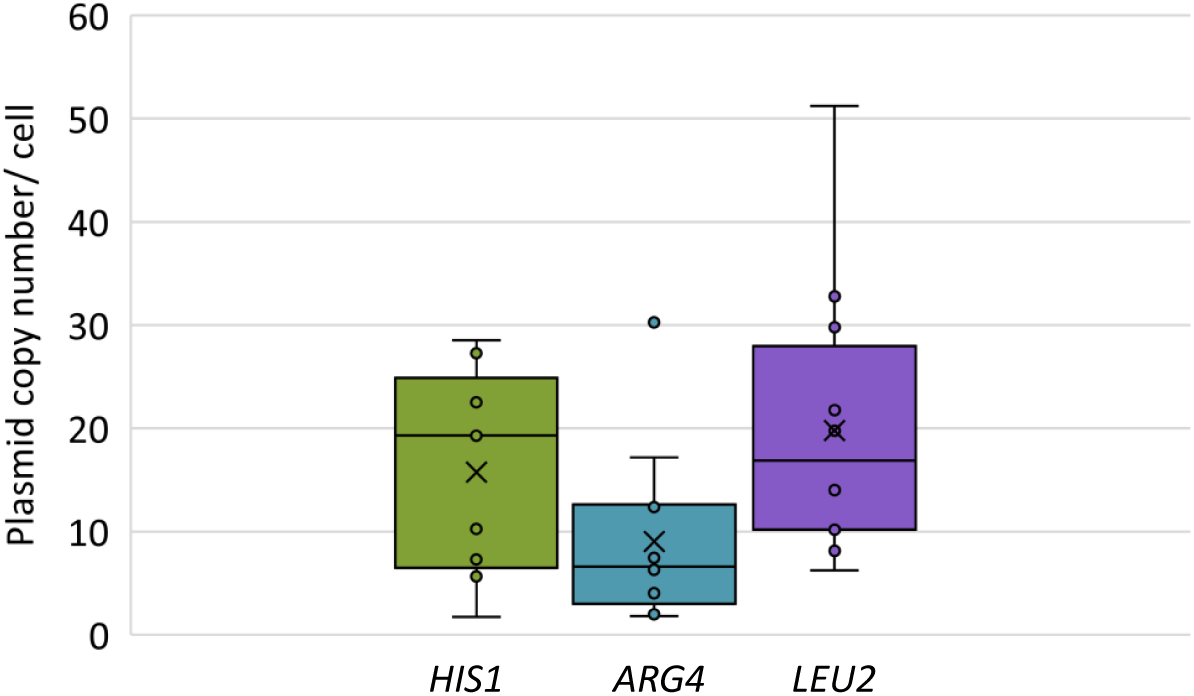
**(A)** Schematic of construction of linear plasmid using primers with telomeric repeats at their ends. **(B)** Comparison of linear plasmids carrying different selection markers and *ORI410* with 0X, 1.5X and 2.5X telomere repeats at its ends: transformation efficiency, proportion of different types of transformants and the number of autonomous transformants. Different markers are represented by different colors and different types of transformants are represented by varying shades of a color (lightest shade representing transients, intermediate shade representing ARS-transformants and darkest shade representing integrants). **(C)** Mitotic stability (%) of ARS-transformants obtained with different linear plasmids with 0X, 1.5X and 2.5X telomere repeats. **(D)** A box plot representing copy number variations of linear plasmids with *CaHIS1*, *CdARG4* or *CmLEU2*, and *ORI410*. The data represents the average copy number of nine independent ARS-transformants (accounting for mitotic stability). In the box plot, dots represent different samples, cross represents mean value and horizontal line represents the median.

Adding TEL repeats to linear plasmids increased the number of ARS-transformants for both *CdARG4* and *CmLEU2* plasmids, compared to those without TEL repeats (Fig. 2B). By contrast, TEL sequence addition increased the TE only for *CdARG4* plasmids, among the three markers tested. Furthermore, adding TEL repeats increased the MS of ARS-transformants by 2-6-fold (MS ~10-35%) for all three markers. Thus, relative to circular plasmids, linearized plasmids with terminal TEL repeats produced more ARS-transformants with higher MS (Fig. 2B and 2C), and the ARS-transformants displayed shorter lag time and doubling time (Table 1, Fig. S1E and S1F). By contrast, when 1.5X TEL sequence was included on circular plasmids, there was no significant change in any of the transformation parameters relative to the corresponding circular plasmids (Fig. S4A and S4B). Thus, it is likely the linear geometry of the plasmids, along with the inclusion of telomere sequence, that resulted in an increase in ARS-transformants and MS (see discussion). Since we found no obvious advantage to including 2.5X vs 1.5X TEL repeats, we used plasmids linearized with the 1.5X TEL repeats in all subsequent studies.

We next asked if linear plasmids carrying 1.5X TEL repeats were maintained autonomously. The ARS-transformants obtained exhibited moderate MS even after three passages, indicating that they were maintained autonomously over a few generations (Table S2). Southern blot of DNA from a pLin-*CdARG4* ARS-transformant with moderate MS (Fig. S2B) showed a single band with electrophoretic mobility similar to that of the naked linear DNA molecule used for transformation. We also recovered pLin-*CmLEU2*-*ORI410* molecules from ARS-transformants in *E. coli* (Fig. S5), demonstrating autonomous replication *in vivo*. Copy number of the linear plasmids in ARS-transformants, measured by qPCR, ranged widely (~2-50 per cell, accounting for MS) (Fig. 2D).

By contrast, *CaHIS1* transformants with moderate MS (45% and 63%) produced larger plasmid-hybridizing bands on Southern blot indicative of genomic integration (Fig. S2B). Further analysis of these *CaHIS1* integrants indicated gene replacement at the native locus either by double cross-over or gene conversion event (Fig. S3B). This is consistent with the idea that plasmids with homologous marker can yield integrants apart from ARS-transformants.

### Effect of marker gene and *bona fide* ORIs on transformation parameters

We next asked to what extent a *bona fide* ORI sequence (*ORI410*) affected the transformation parameters of linear plasmids with different markers. With both homologous markers, *CaURA3* and *CaHIS1,* there were some integration events, initially more frequent for *CaHIS1,* although when propagated under selection, many of the *CaURA3* plasmids integrated (MS ~80-100% and LR <0.1 per generation, Fig. 3B). Furthermore, for *CaURA3* and *CdARG4,* addition of *ORI410* had no effect on the number of ARS-transformants (Fig. 3A), but improved plasmid stability compared to circular plasmids (Fig. 3B). This suggests that there may be a cryptic, intrinsic origin activity within *CaURA3* and *CdARG4* marker fragments (1.3 and 3.1 kb, respectively) that obviates the use of these markers to monitor the contributions of ORIs to plasmid replication and maintenance (discussed below). Similar results for *CaHIS1* and *CdARG4* were evident in different lab strains (Fig. S6). By contrast, addition of *ORI410* on pLin-*CmLEU2* resulted in ~5-fold increase in TE, ~14-fold increase in ARS-transformants and improved plasmid stability (Fig. 3B) relative to pLin-*CmLEU2* (Fig. 3A), suggesting that *CmLEU2* does not carry intrinsic ARS activity seen on other markers.

**Figure 3.**
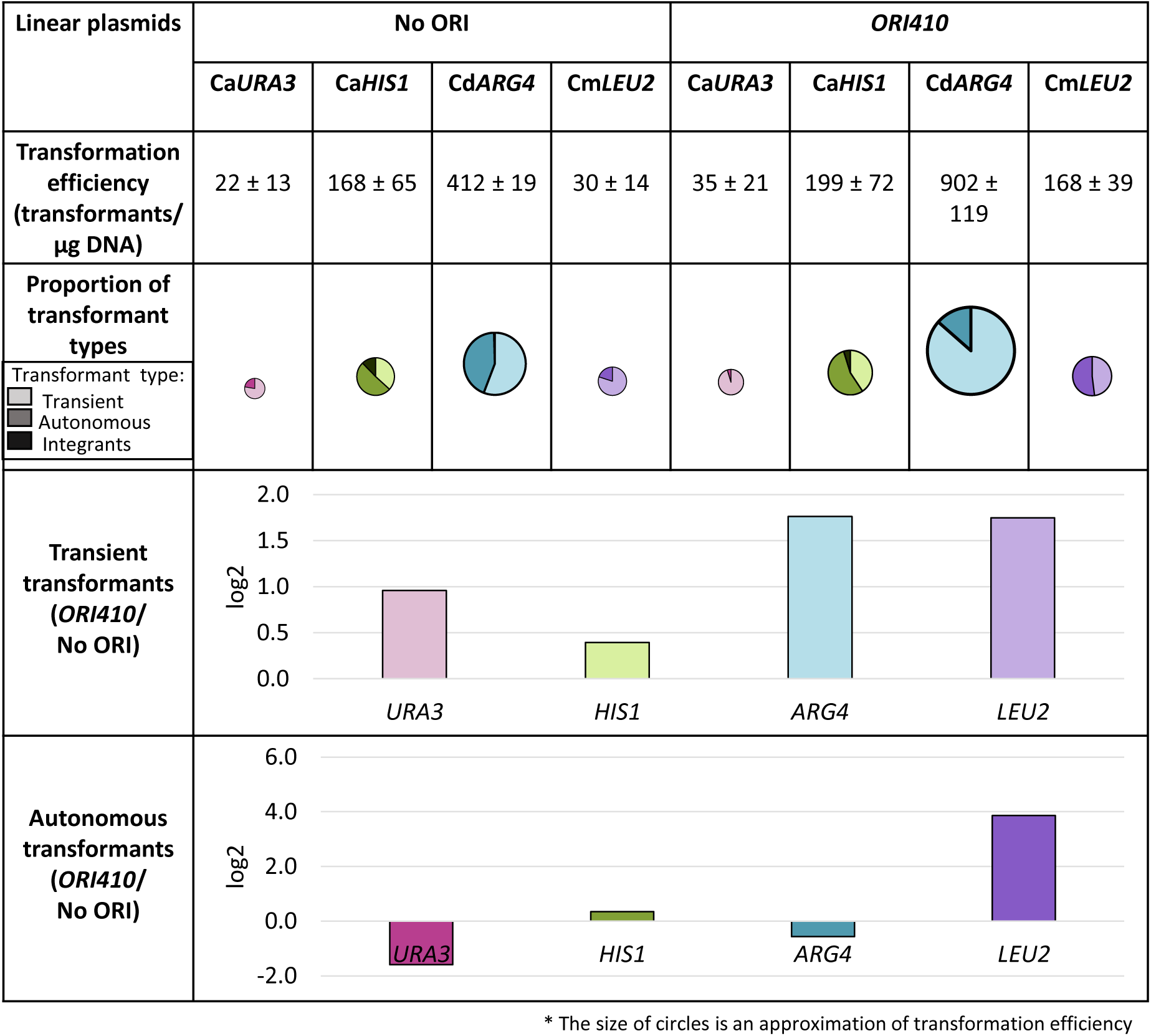
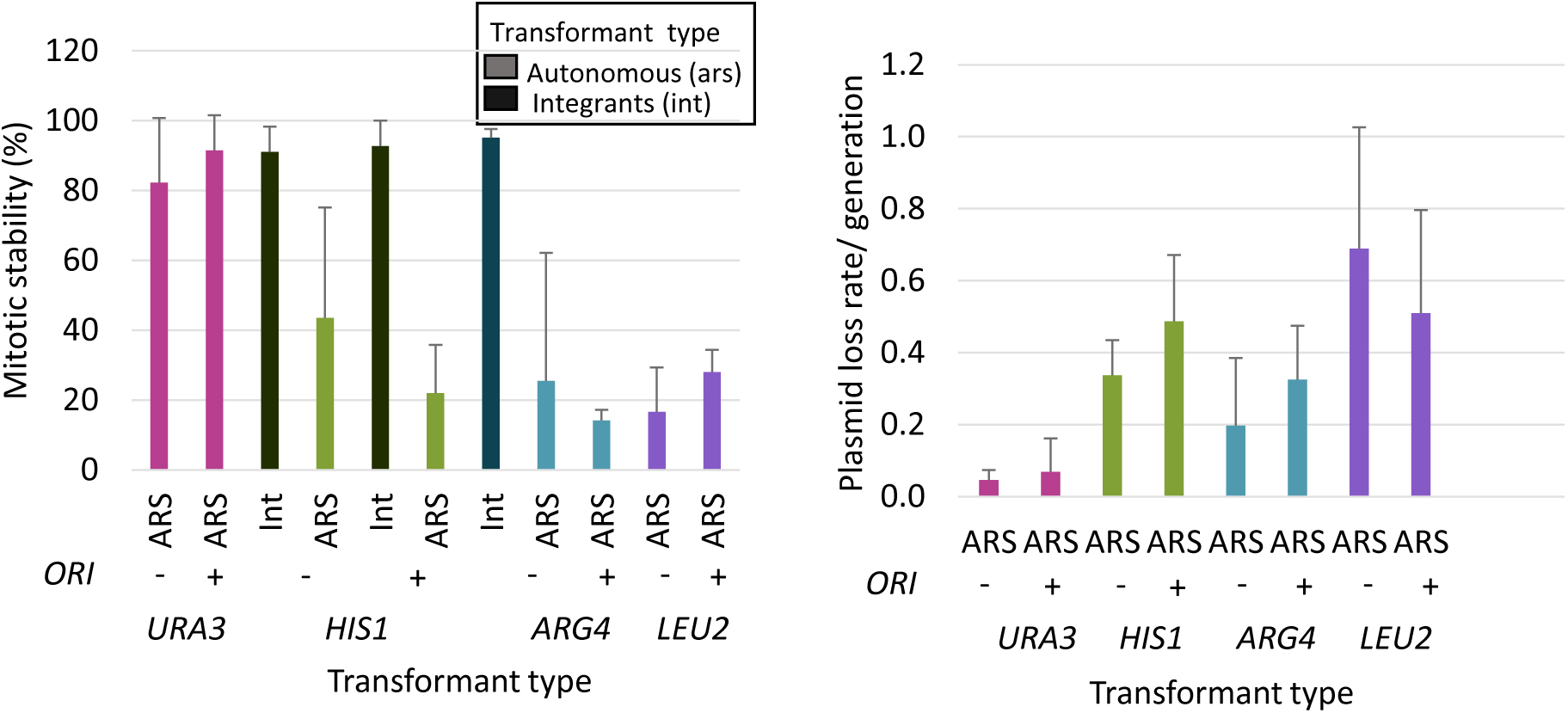
**(A)** Comparison of linear plasmids carrying different selection markers with and without *ORI410*: transformation efficiency, proportion of different types of transformants and log2 value of the ratio of average number of transient or autonomous transformants with ORI to without ORI (*ORI410*/ORI^-^). Different markers are represented by different colors and different types of transformants are represented by varying shades of a color (lightest shade representing transients, intermediate shade representing ARS-transformants and darkest shade representing integrants). The transformation efficiency is an average of three independent experiments. **(B)** Mitotic stability (%) of integrants and ARS-transformants and plasmid loss rate/ generation for ARS-transformants obtained with different linear plasmids with and without *ORI410*. The data represents the average of three independent colonies of each type except for ARS-transformants with *CdARG4* and *CmLEU2* plasmids where it represents the average of six independent colonies. Int: integrants; ARS: ARS-transformants.

Given that all markers were inserted in the same position on a plasmid, which does not have any obvious origin-promoting sequence features, we tested the hypothesis that some feature required for origin firing is present at higher levels in *CaURA3*, *CaHIS1* and *CdARG4* relative to *CmLEU2*; though, many *CaHIS1* ARS-transformants integrate into the genome after additional passages. Since the length of *CaHIS1* and *CmLEU2* are similar, it seems unlikely that marker length is an important factor. Interestingly, the AT content of the two markers *CmLEU2* (62.3%) and *CaHIS1* (63.3%) with higher levels of ORI-dependent ARS-transformants (Fig. 3A), was below the average AT-content of the *C. albicans* genome (66.7%), while the AT-content of *CaURA3* (68.4%) and *CdARG4* (69.1%) was higher than that of the *C. albicans* genome. Thus, it appears that the markers with cryptic ORI function (*CaURA3* and *CdARG4)* that interferes with *bona fide* ORI activity have higher AT content. Of note, neither ORIs alone, nor sequences on markers with possible cryptic ARSs share any obvious conserved primary sequence motifs. Based on its ORI-dependence, *CmLEU2* is the most effective of the markers tested for comparing ORI activity.

### Comparing different *bona fide* origins and *ORI410* fragments

Four ORIs from *C. albicans* (*ORI410* as well as *ORI1055*, *ORI1046* and*ORI246)*, defined previously as ‘*bona fide’* ORIs (28), were inserted into pLin-*CmLEU2* to examine their function compared to no origin. All four *bona fide* ORIs, yielded high TEs (~150-600/µg DNA), ARS-transformants (~75-300/µg DNA) (Fig. 4A) with moderate MS (10-45%) and plasmid LR (0.2-0.7 per generation) (Fig. 4B). *ORI1046* consistently yielded the highest TE and ARS-transformants (~300/µg DNA). Both negative control plasmids pLin-*CmLEU2* (No ORI) or with pro*ORI1088,* a genomic ORC binding region that did not produce replication bubble arcs in 2-D gels (28) gave much lower TE and ARS-transformants (31 and 44/µg DNA). Thus, all four *bona fide* ORIs can drive the origin-dependent autonomous replication of pLin-*CmLEU2* (Fig. 4A).

In *S. cerevisiae*, where the ACS is 11 bp, ARS function is only detected when the transforming fragment is ~100 bp including ACS (7, 8, 37, 38). We asked if two small overlapping fragments (178 bp and 97 bp) derived from *ORI410* (1.2 kb) (28) were able to retain minimum ARS function in *C. albicans*. *ORI410_97* had 2-3 times higher TE (~170 transformants/µg DNA) and ~3 times higher ARS-transformants than no ORI control (Fig. 4A). While the TE and the number of ARS-transformants for *ORI410_97* was lower than for the entire *ORI410*, the ARS-transformants had moderate MS (10-15%) and LR (~0.4 per generation) (Fig. 4B). Thus, an ORI fragment of only ~100 bp can drive linear plasmid replication, and can yield ARS-transformants, which are 2-3-fold more stable than analogous circular plasmid carrying the entire *ORI410* fragment.

**Figure 4.**
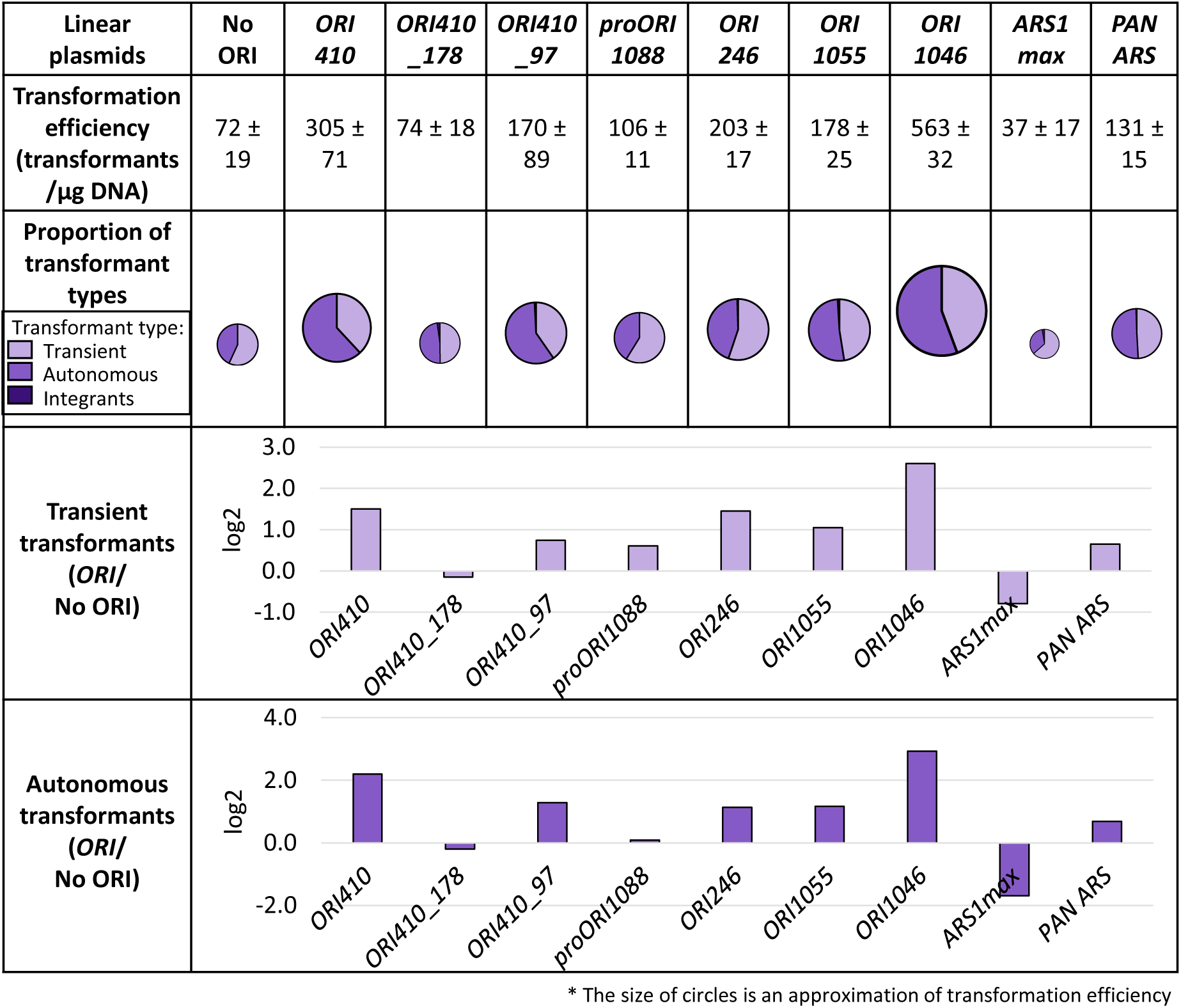
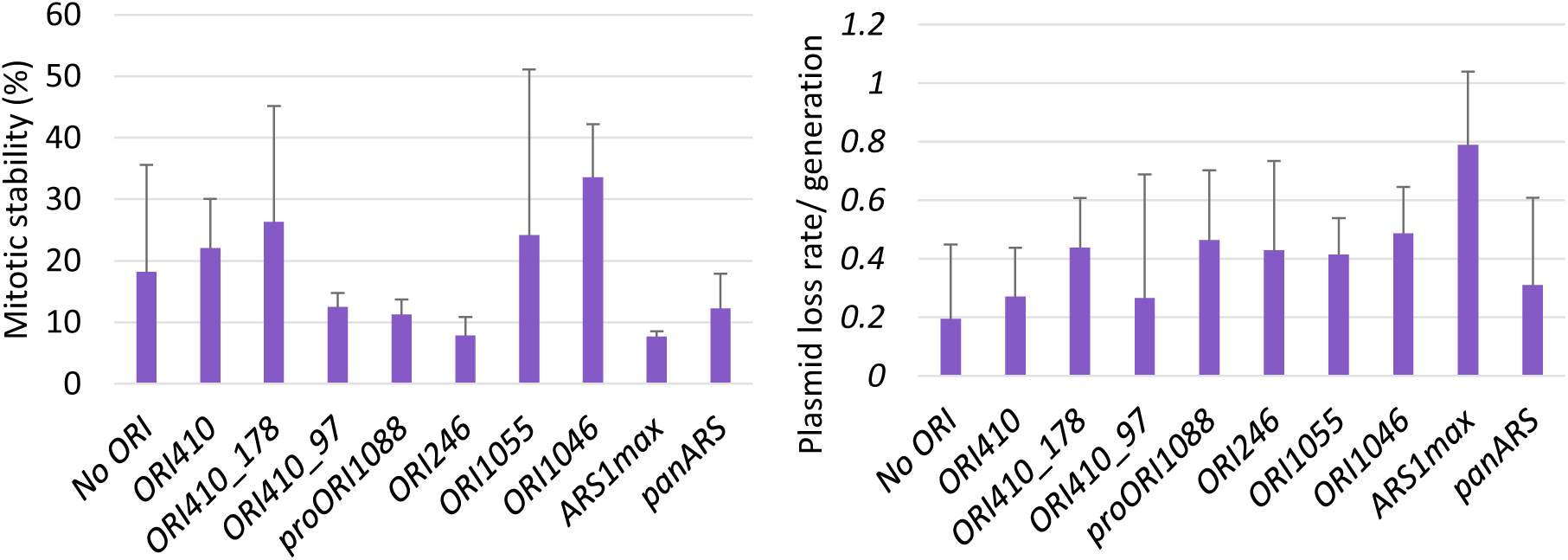
**(A)** Comparison of linear plasmids carrying *CmLEU2* marker with *ORI410* fragments, different *bona fide* ORIs and heterologous ORIs: transformation efficiency, proportion of different types of transformants and log2 value of the ratio of average number of transient or autonomous transformants with ORI to without ORI (*ORI*/ORI^-^). Different types of transformants are represented by varying shades of a color (lightest shade representing transients, intermediate shade representing ARS-transformants and darkest shade representing integrants). The transformation efficiency is an average of three independent experiments. **(B)** Mitotic stability (%) and plasmid loss rate/ generation for ARS-transformants obtained with different linear plasmids mentioned in (A). The data represents the average of six independent ARS-transformants of each plasmid.

### Heterologous ARS sequences

*C. albicans* centromeres are regional and epigenetic, which contrasts with the point centromeres of *S. cerevisiae* (16). Since plasmid replication and origin function were difficult to demonstrate in *C. albicans*, we asked whether heterologous ARS fragments would function in *C. albicans*. The “*panARS*”, a 452bp fragment from *K*. *lactis* genome functions as an active ORI in a range of *Saccharomycotina* yeast species with diverse ARS requirements; in some cases, even more efficiently than average homologous ARSs (e.g. *P*. *pastoris*) (31).

The “*ARS1max*”, an origin from *S. cerevisiae* was selected to drive better growth rates and lower plasmid LR than the original *ARS1* (39). Thus, we tested the ability of both the sequences to direct *C. albicans* plasmid replication on pLin-*CmLEU2* (pLin-*CmLEU2*+*panARS* and pLin-*CmLEU2*+*ARS1max)* relative to pLin-*CmLEU2*+*ORI410* and pLin-*CmLEU2*.

The pLin-*CmLEU2*+*panARS* plasmid resulted in ~2-3-times higher TE and ARS-transformants relative to pLin-*CmLEU2* (Fig. 4A). The *panARS* ARS-transformants had moderate MS (~10-20%) and LR (~0.3 per generation) comparable to those with *ORI410_97* (Fig. 4B). This suggests that the sequence requirements of *C. albicans* origin function are at least partially conserved with those of *K. lactis* among other *Saccharomycotina* species. By contrast, pLin-*CmLEU2*+*ARS1max,* had transformation parameters inferior to those of control plasmid pLin-*CmLEU2;* lower TE, ARS-transformants with lowest MS (<8%) and highest LRs (~0.8 per generation) (Fig. 4A and B) detected for any linear plasmid. This supports the idea that sequence requirements for origins in *C. albicans* (and other yeasts, for example, *Pichia pastoris* (40)) are distinct from those in *S. cerevisiae*.

## Discussion

Early studies seeking potential origin sequences based on their ability to confer high TE, usually used circular plasmids with *CaURA3* as the selectable (and counter-selectable) marker (22, 23, 25, 27). However, most transformants were highly unstable or rapidly integrated into the genome and thus were not useful for autonomous plasmid maintenance. Here we systematically compared four selectable markers in plasmids with circular or linear geometries to monitor the function of ORI sequences. Importantly, a ~100 bp fragment (28) or the heterologous *panARS* (31) was sufficient to provide ARS function on a plasmid in *C. albicans.* This implies that sequence requirements for origin function in *C. albicans* are shared with distantly related yeasts. Nevertheless, the ability of cryptic ARSs on marker sequences to generate transient transformants implies that the sequence requirements for ARS function (and most likely chromosomal ORI function as well) are dependent on sequence context, possibly AT-richness, and other features that are not yet well understood.

An important insight from this work is that **transformant colony size** provides a useful preliminary indicator of plasmid mitotic stability. Presumably, colony size reflects the degree to which the plasmid replication and/or segregation enables growth of individual cells in a population under selective conditions. Specifically, in tiny or ‘pin-point’ colonies (25), plasmids are lost rapidly; in large colonies, plasmids are integrated stably (Table 1). In medium colonies, plasmids are moderately stable (Table 1) because they replicate autonomously, with some cells retaining the plasmid and others losing it. Of note, ARS-transformants with plasmids carrying homologous markers, sometimes integrate in subsequent passages (generating larger colony sub-clones), a property less prevalent with the heterologous markers. Nonetheless, all markers on linear plasmids yield ARS-transformants, which initially can be identified based on colony size.

### Circular vs linear plasmids

In *S. cerevisiae,* linear plasmids and mini-chromosomes were used to study chromosome components and to propagate large segments of DNA (41, 42). However, most work was done with circular plasmids that are readily propagated in *E. coli*; propagation of linearizable plasmids with inverted telomere repeats (29, 43–45) was labor intensive and subject to recombination of the repeats in *E. coli*. Here, a simple approach obviates many of these technical challenges by synthesizing linear plasmids from circles immediately prior to transformation (Fig. 2A). Thus, the two plasmid geometries are directly comparable, differing only by the presence or absence of 1.5X TEL repeats.

Does the presence of TEL sequence alone improve the segregation of linear vs circular plasmids? In *C. albicans* as in *S. cerevisiae,* adding TEL sequences to linear plasmids improves their stability (46) (Fig. 2C). However, adding TEL sequences does not improve circular plasmid segregation in *C. albicans* (Fig. S4); by contrast, TEL sequences on circles stabilized *ScARS* plasmids (47) and antagonized the segregation of *ScCEN* plasmids (48). Thus, *Ca*TEL sequence function is required for autonomous linear plasmid maintenance and is dependent upon its geometry: in a chromosome end context, but not within a circular context. This supports the idea that interactions between non-terminal TEL DNA and telomeric proteins likely differ between *C. albicans* and *S. cerevisiae* and that linear plasmids require telomere ends to remain stable.

In *S. cerevisiae*, non-centromeric plasmids are retained in the mother cells due to their attachment with nuclear membrane (49) as well the presence of a diffusion barrier at the bud neck (50). It is tempting to speculate if this is also true for circular plasmids (with or without TEL) in *C. albicans*. Whether and how the linear plasmids might be more able to segregate to daughter cells remains to be explored.

### Effect of selectable markers

Comparison of the markers found that *CaURA3* was not ideal, which explains difficulties in many earlier investigations (25, 26) and addition of *LEU2 or HIS1* to *CaURA3* plasmids relied on *URA3* selection as well (22, 27). Studies selecting for *IMH3^R^* or *CaADE2* found that putative ARS-transformants integrated at high frequency (21, 25). Sometimes integration events involved and/or altered the putative origin structure (24) and the resulting plasmids were not maintained autonomously. Notably, *CaURA3* linear plasmid produced very few ARS-transformants, with or without ORI addition, and these eventually integrated into the genome (Fig. 3). By contrast, ARS-transformants with either *CaHIS1, CdARG4 or CmLEU2* were maintained over three passages (Table S2).

We suggest the appearance of transient transformants cannot be used to define origin function on a plasmid, especially when *CaURA3* marker with latent origin activity is used. Therefore, transients seen with *ORI7-L1* and *–R1* cannot be used to make conclusions about the function of these chromosomal regions as origins, especially since the published data lacks a control plasmid containing the *CaURA3* marker without an origin (26). Transient transformants with these origins have been used to postulate that centromere function required a pre-existing origin. However, our results showing that these chromosomal regions do not act as origins, together with published neocentromere locations at chromosomal regions that did not contain pre-existing origins (51, 52), support a model where kinetochore assembly can convert a non-origin region to an origin. Furthermore, if many genome sequences can recruit replication factors and provide weak origin function on a plasmid as in *S. pombe* (14, 53), it is not surprising that sequences within neocentromere regions may recruit origins to new loci. The dramatically increased origin efficiency of the neocentromeric loci is likely due to neocentromere-mediated recruitment of replication initiation activities like Cdc7-Dbf4, which is normally found at wild-type centromeres (54).

Heterologous *CdARG4* did not integrate frequently, yet, it gave high numbers of ARS-transformants in the absence of an added ORI. We posit that both *CaURA3* and *CdARG4* have weak intrinsic ARS activity and that this may compete with a *bona fide* ORI when both are on a plasmid. This suggests that *C. albicans,* like *S. pombe,* has “cryptic origins” (55), i.e., sites that are normally not used for replication initiation, yet have the potential to form active replication origins. It also suggests that, once a cryptic origin has been established, it can continue to function, perhaps because, once well-established in an ARS-transformant, a weak origin may be more likely to fire in the next cell cycle. What the requirements for cryptic ARS function are, remains elusive. We cannot rule out the possibility that chromatin structure and topological constraints might affect ARS activity.

Why might inefficient ORIs interfere with *bona fide* ORI activity? In *S. cerevisiae*, two ORIs in close proximity in the genome interfere with each other (56). Three mechanisms were proposed to explain this: 1) timing of ORI firing might differ such that the non-firing ORI is replicated passively; 2) DNA at the two ORIs might interfere topologically (e.g., via altered supercoiling); or 3) the two origins may compete for a limited number of licensing factors (e.g., ORC-associated proteins) (56). Interestingly, the orientation of ORC sites relative to one another could also be relevant (57) and all six predicted ORC sites (28) on *CmLEU2* are oriented in the same direction, while predicted ORC sites (28) on the other three markers were found in both orientations. While mechanisms of *Ca*ORI and *Sc*ORI firing are likely to differ to some degree, these options may explain the phenomenon in *C. albicans* as well.

Most organisms do not have highly defined ARS consensus sequences, and appears that this is the case in *C. albicans* as well. In *S. pombe*, ORIs have average AT content ranging from 72-75% (58), with an average of 64% in the genome. *CaURA3* and *CdARG4* have 68.4% and 69.1%, respectively, with an average of 66.7% AT content in the genome. Furthermore, for all four markers on linear plasmids, the number of polyA tracts (≥3 nucleotides, normalized for marker length) correlated well (R^2^= 0.85) with the number of transients obtained. This is consistent with the idea that AT rich sequences and/or polyA tracts may attract replication factors, and acquire cryptic ORI function. This, in turn, might interfere with *bona fide* ORI firing on the plasmid by mechanisms like those proposed for *S. cerevisiae* (56).

### Testing origin function

The linear *CmLEU2* plasmid backbone provided the first opportunity to compare the efficiency of different *bona fide* ORIs, *ORI410*-derived fragments (28) as well as heterologous origins (31, 39). All four *bona fide* ORIs yielded high numbers of ARS-transformants as well as moderate MS and LRs (Fig. 4). We do not know why the 178 bp fragment, *ORI410_178,* had little or no obvious origin function while a smaller fragment derived from it (*ORI410_97)* was active. Clearly, DNA primary sequence is not sufficient to confer ORI function. We presume that sequence features, together with their context relative to other plasmid components affect ORI activity, which has been seen on *Sc*ARS plasmids as well (59). Importantly, the synthetic *panARS*, which was derived from *K. lactis* (31), had transformation parameters similar to those of *ORI410_97.* Thus, the requirements for replication origin function *in C. albicans* are at least partially conserved with other *Saccharomycotina* species and *panARS* provides a heterologous ORI that should not integrate into the genome. We suggest that it might be possible to whittle down the 452 bp *panARS* to generate a relatively good heterologous ORI of ~100 bp.

### Summary

ARS function can be studied in *C. albicans* using a heterologous marker and a *bona fide* ORI of as small as ~100 bp or the heterologous *panARS* on linear plasmids carrying 1.5X TEL ends. Importantly, linear plasmid conformation greatly facilitates transformation efficiency and mitotic stability. Unexpectedly, the choice of selectable marker has a major effect on the degree to which plasmids are maintained autonomously. To date, *CmLEU2* is the single marker that has a low level of intrinsic cryptic origin activity, and rare integration events, making it ideal for studying origin activity on a plasmid. The linear plasmids described here fill a major gap in the tools available for conventional molecular manipulations of *C. albicans* and will facilitate our ability to study molecular aspects of ORI, telomeric, and centromeric structure and function.

## Materials and Methods

### Strains, plasmids, primers and growth conditions

Yeast strains and plasmids used are listed in Table 2. Primers used are provided in Table 3. *C. albicans* strains were grown at 30 °C in YPAD medium (60) or SD minimal medium or SD-Complete medium (60) containing leucine at 170 mg/l and all other amino acids (Sigma Aldrich, USA) at 85 mg/l.

**Table 2.** **(A)** List of strains used in the study

| Strain no. | *C. albicans* strains | Genotype | Source | Used with |
| --- | --- | --- | --- | --- |
| YJB-T 45 | BWP17 | *ura3::λimm<sup>434</sup>/ura3::λimm<sup>434</sup>*<br>*his1::hisG/his1::hisG*<br>*arg4::hisG/arg4::hisG* | (65) | CaURA3 |
| YJB-T 72 | SN76 | *ura3-iro1::imm<sup>434</sup>/ura3-iro1::imm<sup>434</sup>*<br>*his1::hisG/his1::hisG* *arg4Δ/arg4Δ* | (30) | CaHIS1,<br>CdARG4 |
| YJB-T 736 | SN152 | *ura3-iro1::imm<sup>434</sup>/URA3-IRO1*<br>*his1::hisG/his1::hisG* *arg4Δ/arg4Δ,*<br>*leu2Δ/leu2Δ* | (66) | CmLEU2 |
| YJB-T 65 | SN95 | *ura3-iro1::imm<sup>434</sup>/URA3-IRO1*<br>*his1::hisG/his1::hisG* *arg4Δ/arg4Δ* | (30) | CaHIS1,<br>CdARG4 |

**(B)** List of plasmids used in the study
| Plasmid no. | Description | Source |
| --- | --- | --- |
| BJB-T1 | pGEM-*URA3* | (65) |
| BJB-T226 | pGEM-*URA3-ORI410* | This study |
| BJB-T2 | pGEM-*HIS1* | (65) |
| BJB-T140 | pGEM-*HIS1-ORI410* | This study |
| BJB-T391 | pGEM-*CdARG4* | This study |
| BJB-T234 | pGEM-*CdARG4-ORI410* | This study |
| BJB-T230 | pGEM-*CmLEU2* | This study |
| BJB-T231 | pGEM-*CmLEU2-ORI410* | This study |
| BJB-T398 | pGEM-*CmLEU2-ORI410_178* | This study |
| BJB-T399 | pGEM-*CmLEU2-ORI410_97* | This study |
| BJB-T400 | pGEM-*CmLEU2-proORI1088* | This study |
| BJB-T401 | pGEM-*CmLEU2-ORI246* | This study |
| BJB-T402 | pGEM-*CmLEU2-ORI1055* | This study |
| BJB-T403 | pGEM-*CmLEU2-ORI1046* | This study |
| BJB-T404 | pGEM-*CmLEU2-ARS1max* | This study |
| BJB-T405 | pGEM-*CmLEU2-panARS* | This study |
| BJB-T227 | pGEM-*URA3-ORI7-R1* | This study |
| BJB-T228 | pGEM-*URA3-ORI7-L1* | This study |

**Table 3.**
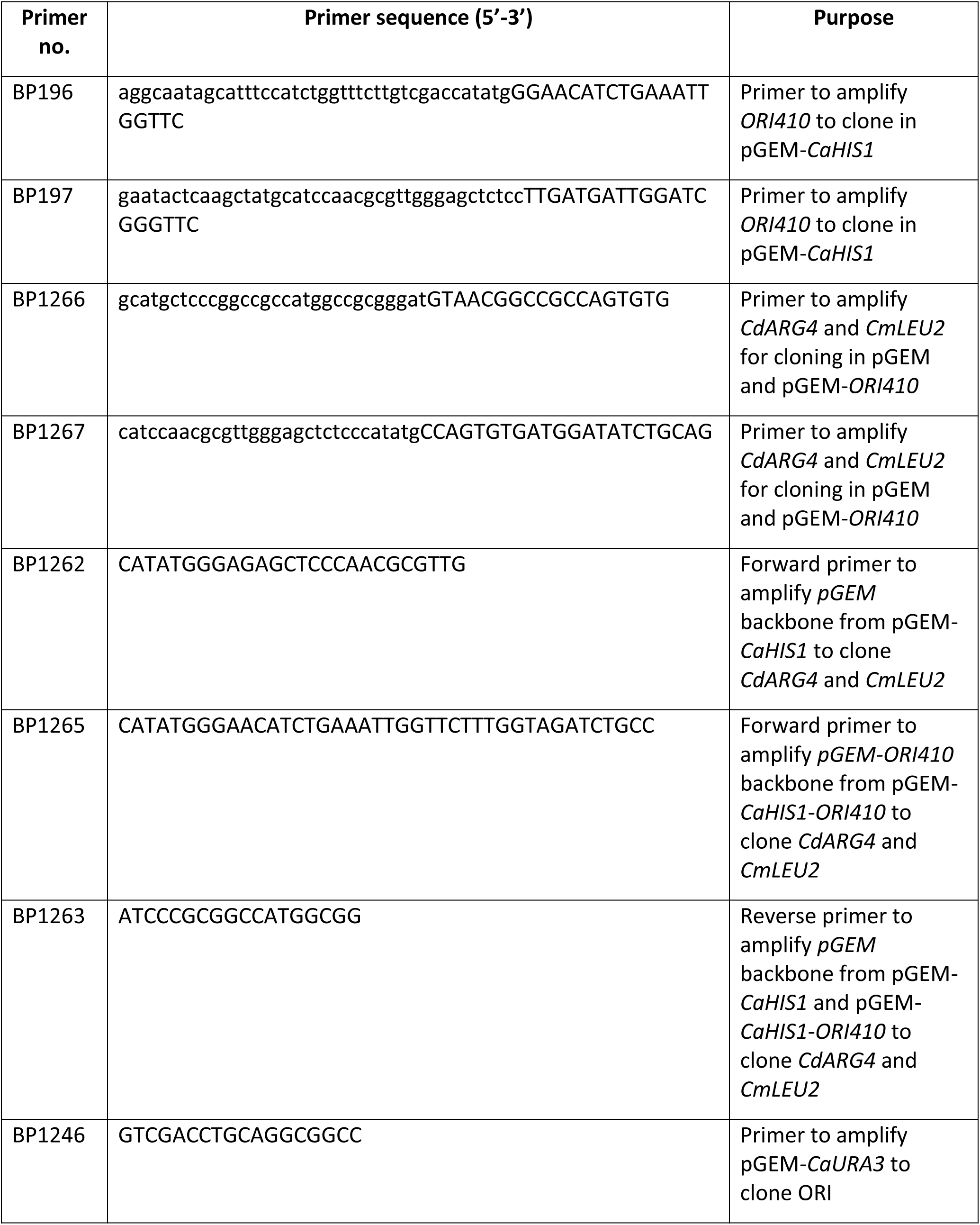
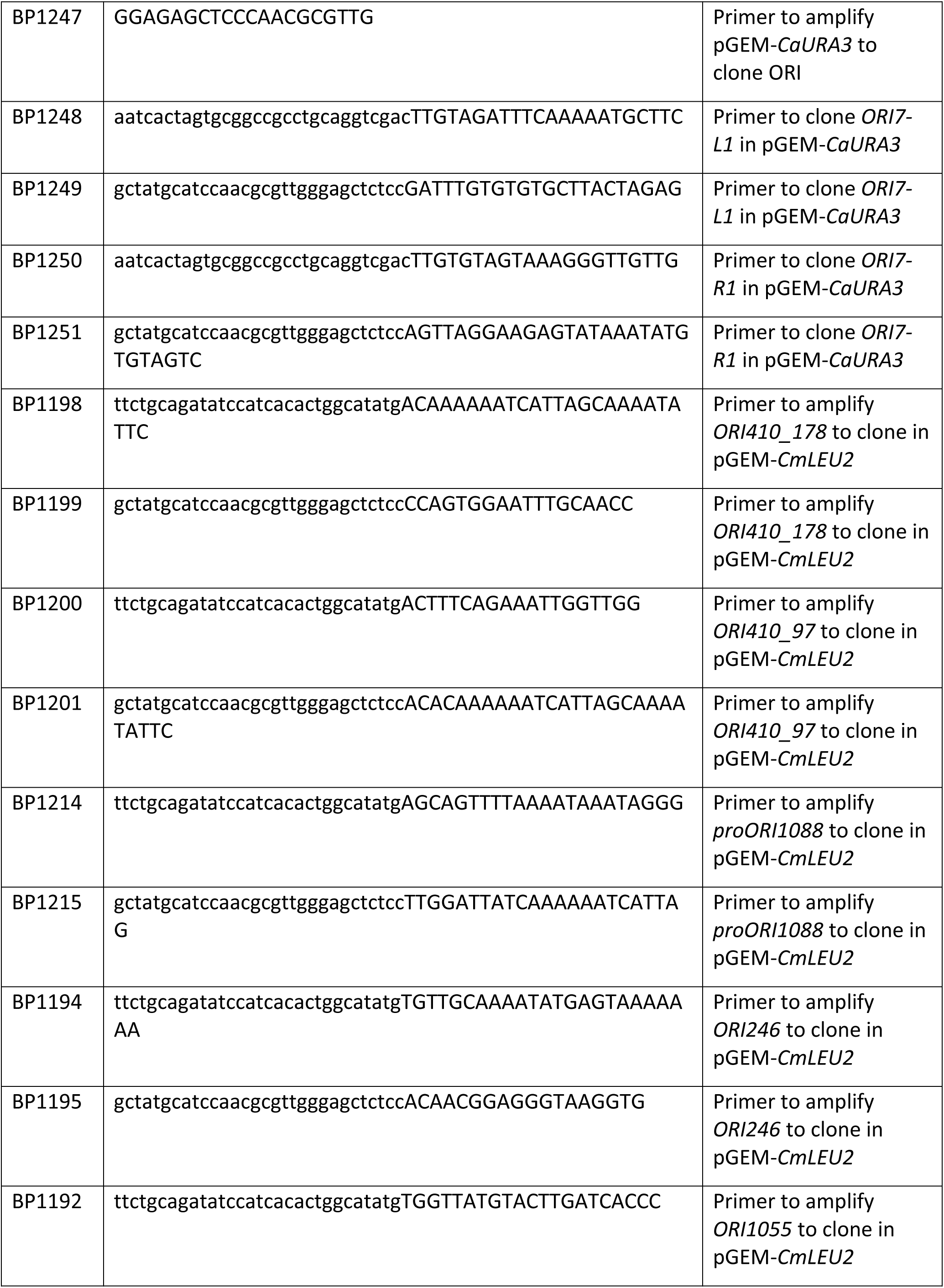
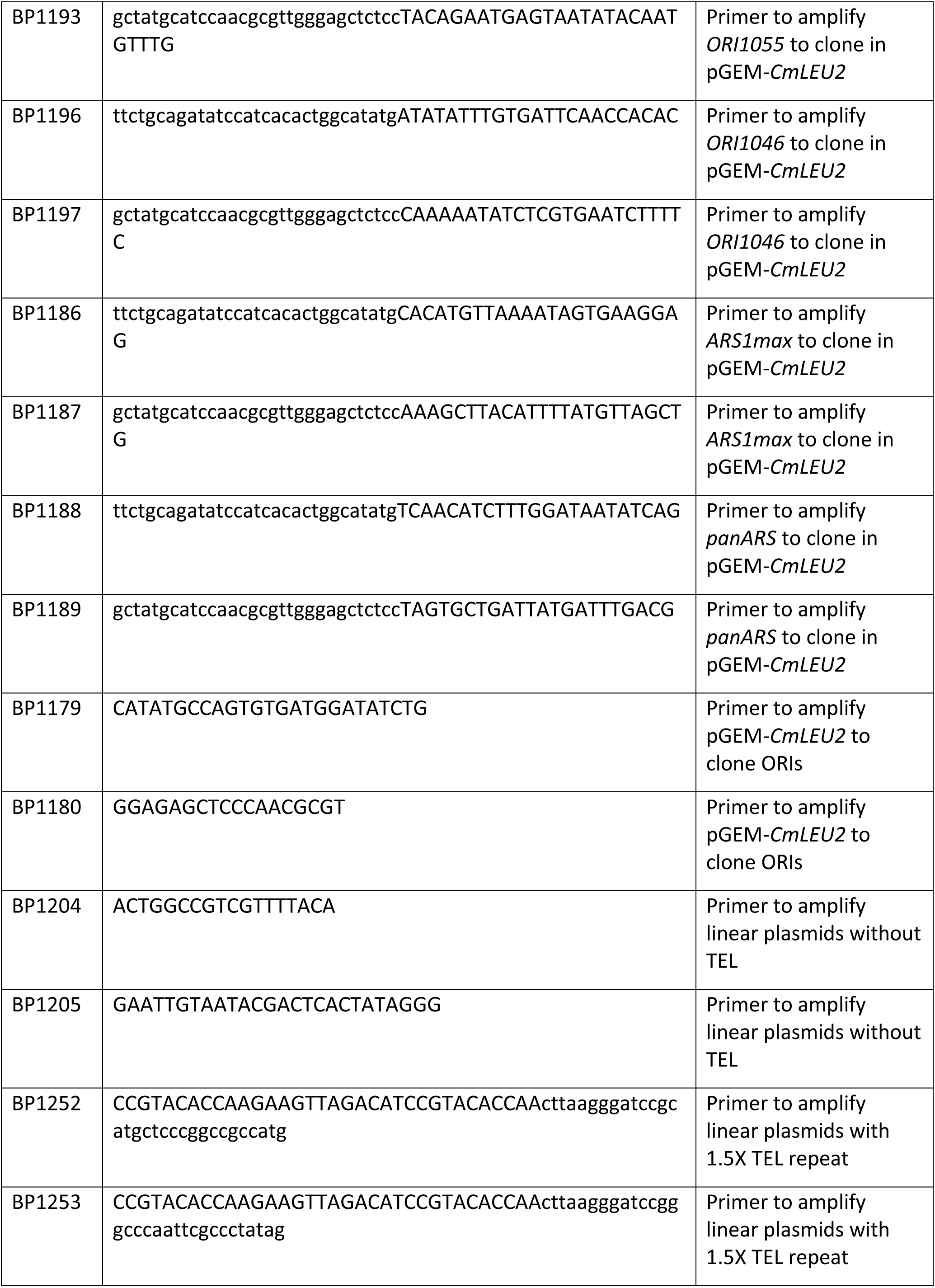
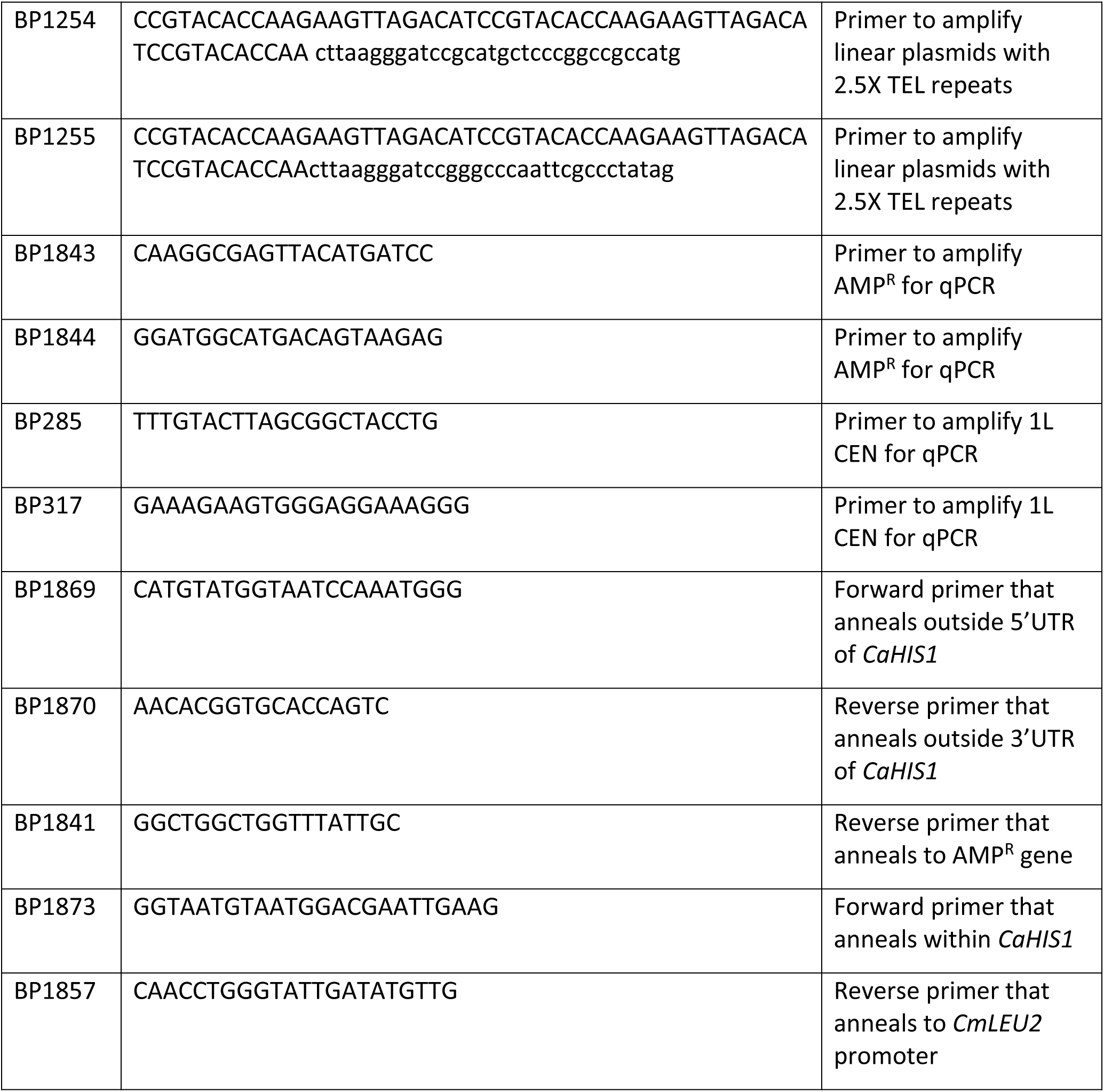
List of primers used in the study

*E. coli* DH5α was used for all cloning experiments and was grown in LB medium (60) at 37°C with Ampicillin (Sigma Aldrich, USA) at 100 µg/ml.

### Cloning of selection markers and ORIs in plasmids

Selection markers and ORIs were amplified with primers (Table 2) carrying 15-40 bp homology to the vector and ~20 bp homology to the marker or ORI fragment. Amplified vector and insert (1:3 ratio) were assembled in 20 µl Gibson reaction (61) as per manufacturer’s instructions (NEB, USA) and 2 µl was transformed into chemically competent *E. coli* (NEB, USA). Following selection on LB+ampicillin overnight, recombinants were detected by colony PCR using primers to the vector, outside of the cloning sites. Putative positive clones were then confirmed by Sanger sequencing.

### Construction of linear plasmids

Linearizing primers (BP1252, BP1253, BP1254, BP1255; Table 2) contained (from 5’ and 3’) 34 or 57 nt telomere sequence (36); *Afl*II and *Bam*HI recognition sites; and then homology to the plasmid *Aat*II site. Linear plasmids were amplified from circular plasmids (Fig. 2A) by two-step PCR using Kappa HiFi HotStart polymerase (Roche, Swtizerland). Cycling conditions were: 98°C denaturation step for 30 sec; 30 cycles of 98°C (10 sec), 72°C (30 sec/kb); final extension, 72°C for 10 min.

To generate linear plasmids without telomeric ends, the circular plasmids were amplified with primers (BP1204, BP1205; Table 2) using Phusion polymerase (Thermo Fisher Scientific, USA). Cycling conditions were: 98°C denaturation step for 30 sec; 25 cycles of 98°C (10 sec), 60°C (30 sec), 72°C for (30 sec/kb); final extension step at 72°C for 10 min.

### Colony PCR

A small portion of the colony was resuspended in the PCR reaction with *Taq* polymerase (Hy-Taq Ready Mix, Hy-labs, Israel). Cycling conditions were: 95°C denaturation step for 5 min; 25 cycles of 95°C (30 sec), annealing at a primer-dependent temperature (30 sec), 72°C (1 min/kb); final extension step at 72°C for 5 min.

### High efficiency transformation of *C. albicans*

*C. albicans* transformation was carried out as described in (62) with the only difference that DTT was added at a final concentration of 25 mM and after 45 min incubation with LiAc-TE, the cells were further incubated with DTT for 1.5 h.

### Mitotic stability assay

Yeast transformants were inoculated into SDC(-AA) (selective) media and grown overnight at 30°C. The cultures were 10-fold serially diluted and appropriate dilution to yield 100-200 colonies was plated onto both SDC (-AA) and SDC plates. The plates were incubated at 30°C and the number of colonies were counted after 2 days. Mitotic stability was calculated as:

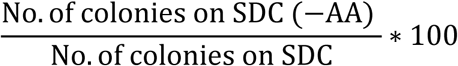

### Plasmid loss assay

Yeast transformants grown overnight in SDC (-AA) for mitotic stability assay were diluted 100-fold into SDC media and grown overnight at 30°C. The cultures were 10-fold serially diluted and 5 µl of each dilution was spotted on both SDC (-AA) and SDC plates. The plates were incubated at 30°C for 2 days and the number of colonies were counted from the highest dilution where they were well separated. The proportion of cells that retained the plasmid without selection was calculated as (No. of colonies on SDC (-AA)/ No. of colonies on SDC) from the same dilution. The plasmid loss rate was then determined as described in Longtine et. al. (47).

### Southern blotting

The genomic DNA was extracted from 10 ml overnight grown cultures in SDC (-AA) as described in (62). 15-20 µg genomic DNA was digested overnight with *Apa*I and run on 1% agarose gel for 16-20 h at 1.4V/cm. Southern blotting was performed as described in (63). PCR fragment of *AMP^R^* gene was used to probe the plasmids on the blot.

### qPCR to determine plasmid copy number

qPCR was carried out with the genomic DNA from autonomous transformants using SYBR green master mix (Bio-Rad, USA) as per manufacturer’s protocol in BioRad CFX96 Touch^TM^ Real-Time PCR Detection System. Cycling conditions were: 95°C (3 min); 40 cycles of 95°C (5 sec), 60°C (30 sec); melt curve from 65.0 to 95.0 for 5 sec. The *AMP^R^* gene was used to determine plasmid copy number and *CEN* of chromosome 1 was used as a reference gene. The two primer sets used had similar efficiency in the reaction; therefore, fold change in the copy number of plasmids was determined relative to the genomic control. Copy number of plasmids was calculated as:

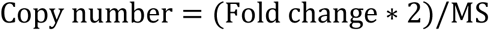

### Growth rate determination

From fresh transformation plate, three independent colonies per colony size were inoculated into 2 ml SDC (-AA) and grown overnight at 30°C, 250 rpm. 50 µl of cell culture was washed with ddH_2_0 and re-suspended in 1 ml SDC (-AA); 10 µl was inoculated in 100 µl SDC (-AA) in a 96-well round bottom sterile polystyrene plates (Corning). For tiny colonies that could not be propagated in liquid media, three independent colonies were directly inoculated from the plate into 100 µl SDC (-AA). The plate was subsequently incubated at 30°C in a Tecan Infinite F200 Pro (Tecan, Switzerland) microplate incubator/spectrometer with a shaking duration of 900 sec and the OD_600_ was collected every 15 min over a 24 h period. OD vs Time was plotted to generate growth curves.

### Recovery of pLin-*CmLEU2*-*ORI410* plasmid from autonomous transformants

The genomic DNA from the yeast transformants was digested with *Bam*HI (NEB, USA) to cut the linear plasmid at both the ends resulting in removal of telomere repeats. The digested DNA was ligated overnight with T4 DNA ligase (Thermo Fisher Scientific, USA). The ligation product was transformed into electrocompetent *E. coli* (64) and the clones obtained were confirmed by PCR primers flanking the ligation site followed by sequencing.

## Acknowledgements

We thank members of the Berman lab for stimulating discussions throughout the work, Anton Levitan for help with sequence analysis and Ella Segal, Andre Maicher, Shay Bramson and Sophia Hirsch for technical assistance. We thank Ella Segal, Laura Burrack and Amnon Koren for helpful comments on the manuscript and for Southern blotting experiments. This work was supported by the Israel Science Foundation (314/13) and by the European Research Council Advanced Award (340087, RAPLODAPT) to J.B., and a PBC postdoctoral fellowship to M.A.T.

